## Supplementary Material for "A cryo-EM structure of KTF1-bound polymerase V transcription elongation complex"

### Supplementary Figures

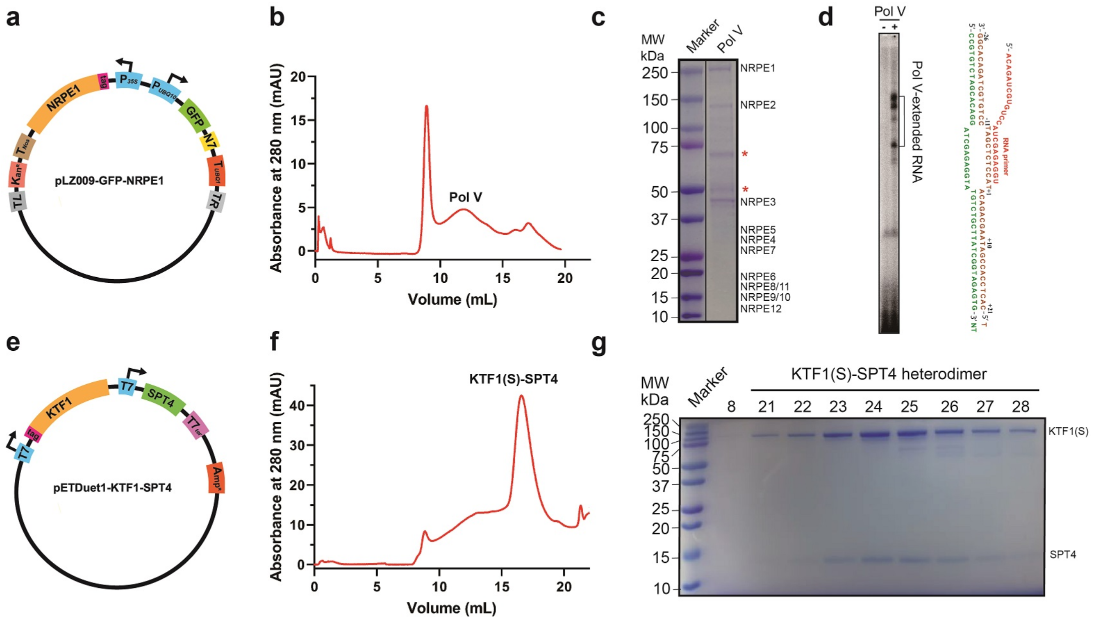


**Supplementary Figure 1. The procedure for preparation of endogeous Pol V and recombinant KTF1-SPT4. (a)** The pLZ009-GFP-NRPE1 construct encodes the NRPE1 with a 3xFLAG-tag at its N-terminal region under control of $P_{35S}$ promoter and $T_{NOS}$ terminator, the GFP gene under control of $P_{UBQ10}$ promoter and $T_{UBQ1}$ terminator, and a kanamycin-resistance gene. TL and TR indicate boundaries of DNA fragment to be integrated into the T87 genome. **(b)** Size exclusion chromatogram of the epitope-tagged *A. thiniana* Pol V. **(c)** The Coomassie Brilliant Blue staining SDS-PAGE result of the purified epitope-tagged *A. thiniana* Pol V. Red stars, impurity. The experiments were repeated 3 times. **(d)** In vitro radiochemical transcription result shows Pol V extends the 23-nt RNA primer. The experiments were repeated 3 times. **(e)** The pETDuet1-KTF1-SPT4 construct encoding His-tagged *A. thiniana* KTF1(1-712) and SPT4. **(f)** Size-exclusion chromatogram of the purified KTF1(S)-SPT4 complex and **(g)** the Coomassie Brilliant Blue staining SDS-PAGE result of the eluted peak fractions for the KTF(1-712)-SPT4 complex. KTF1(S): KTF(1-712). The experiments were repeated 3 times.

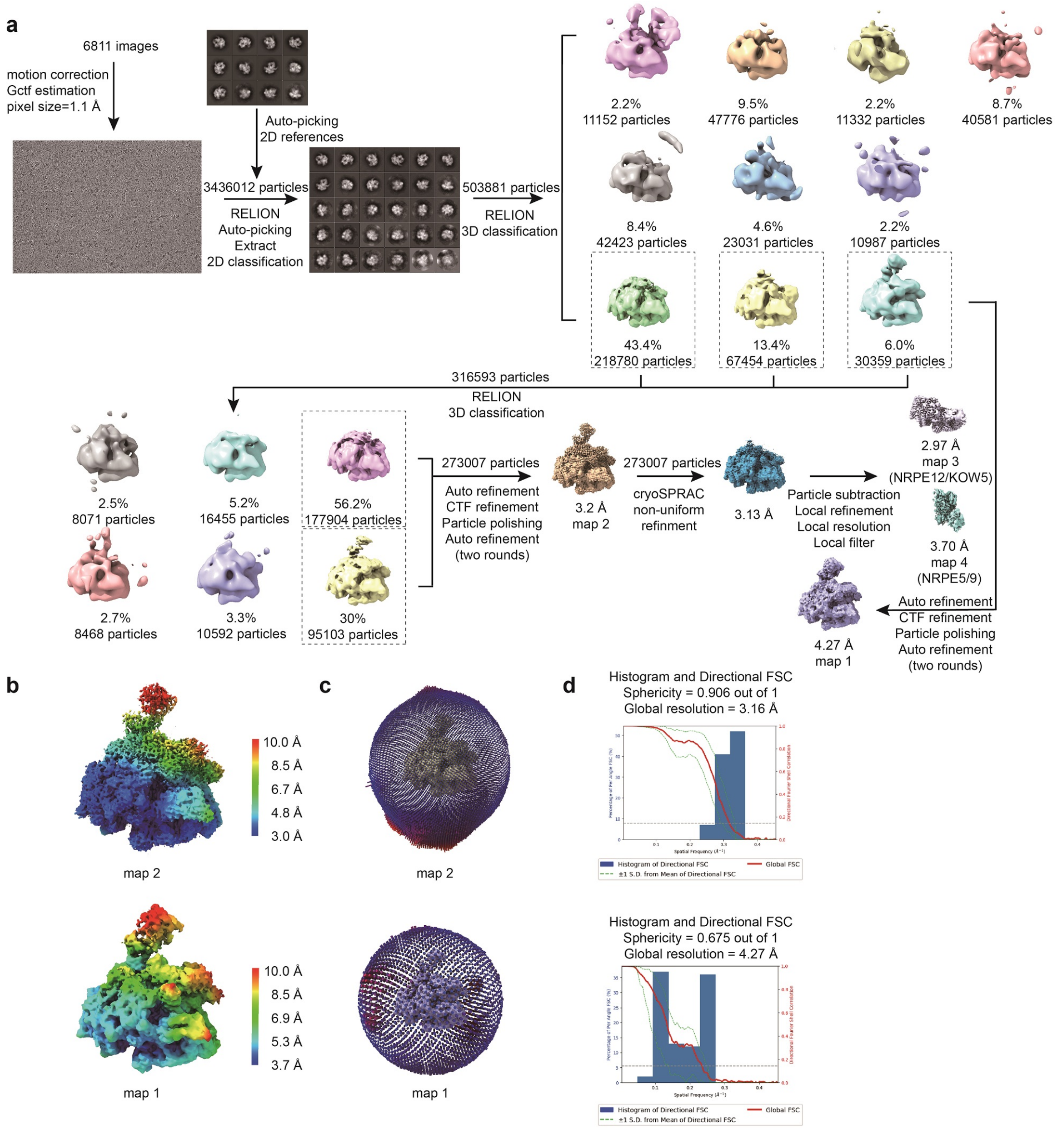


**Supplementary Figure 2. The procedure of cryo-EM map reconstruction of the KTF1-bound Pol V TEC. (a)** The flowchart of data processing of the KTF1-bound Pol V TEC. **(b)** The local resolution of the two resconstructed cryo-EM maps (map 1 and map 2) of the KTF1-bound Pol V TEC. **(c)** The angular distribution of single-particle projections of the two *A. thaliana* KTF1-bound Pol V TEC maps (map 1 and map 2). **(d)** The 3D FSC plot of the two KTF1-bound Pol V TEC maps (map 2 on the upper panel and map 1 on the lower panel).

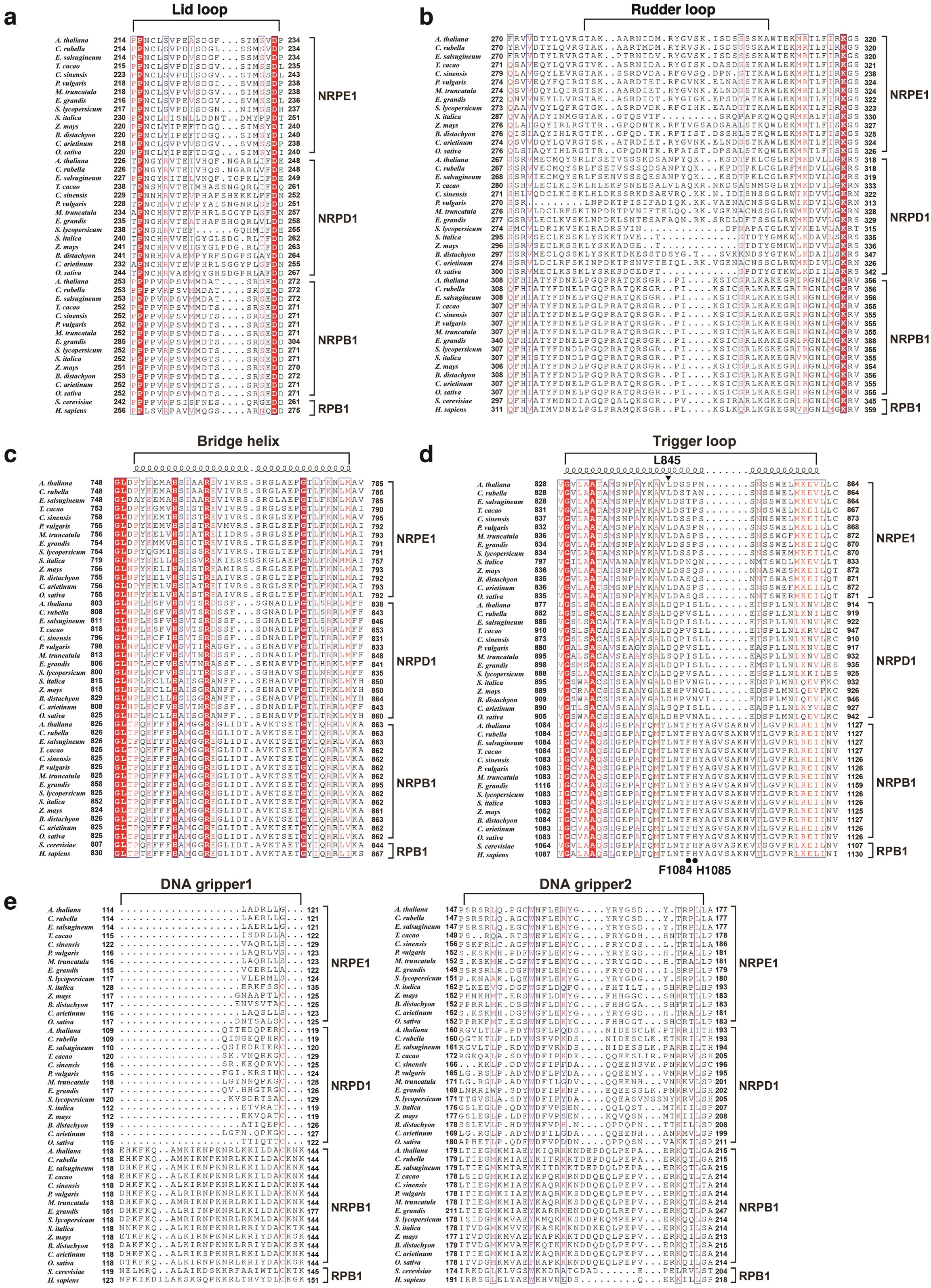


**Supplementary Figure 3. The sequence aligment of (a) lid loop, (b) rudder loop, (c) bridge helix, and (d) trigger loop in the active-site cleft of Pol V, Pol IV, and Pol II.** The residue L845 of A. thiniana Pol V NRPE1 is indicated by filled triangle and the residues F1084 and H1085 of *S. cerevisiae* Pol II RPB1 are indicated by filled circles. **(e)** The sequence aligment of the two dsDNA grippers of Pol II.

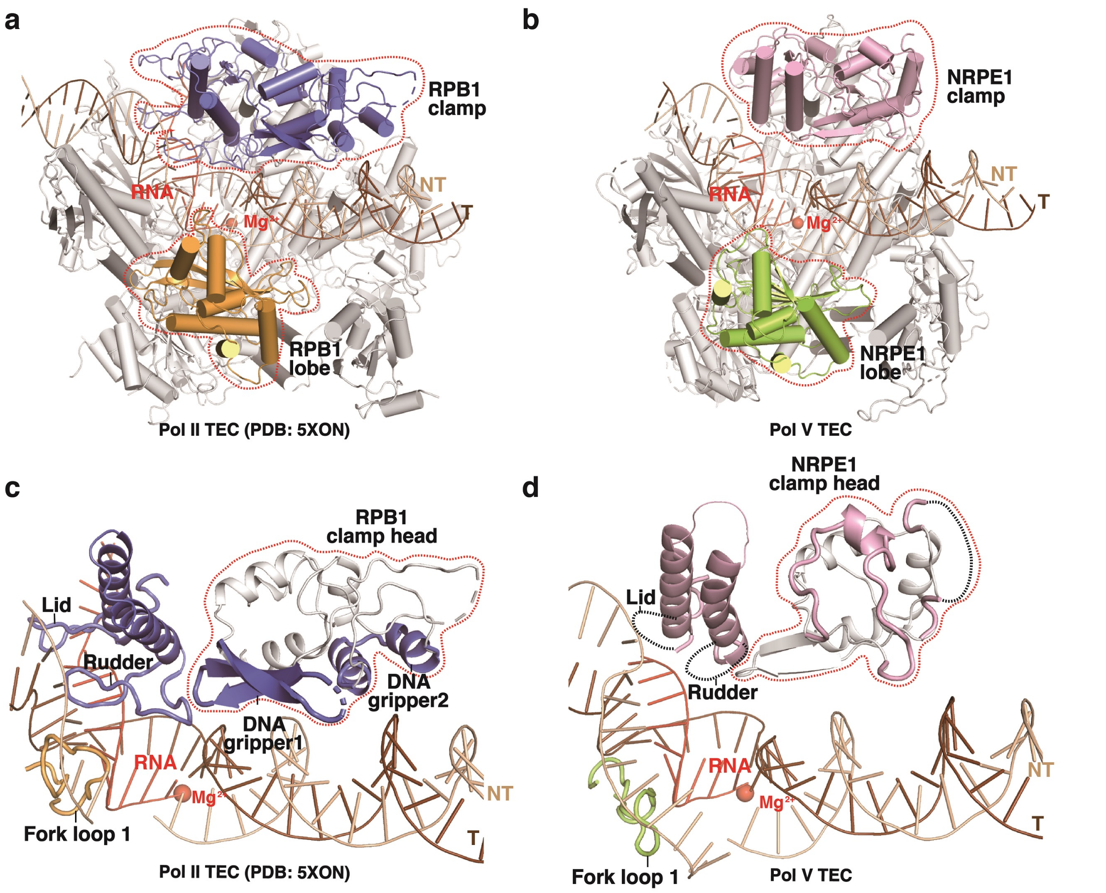


**Supplementary Figure 4. Pol V clamp domain loosely interacts with DNA and RNA. (a)** The cartoon shows the position of the clamp domain and the lobe domain of the Pol II TEC (PDB: 5XON)[1]. The clamp domain of Pol II is colored in slate, and the lobe domain of Pol II is colored in orange. **(b)** The cartoon shows the position of the clamp domain and the lobe domain of the Pol V TEC. The clamp domain of Pol V is colored in pink, and the lobe domain of Pol V is colored in green. **(c)** The cartoon presentation of interactions between Pol II and the DNA/RNA, and **(d)** the interactions between Pol V and the DNA/RNA. Red dashes depicit the clamp head domain of the largest subunit of Po l II or Pol V.

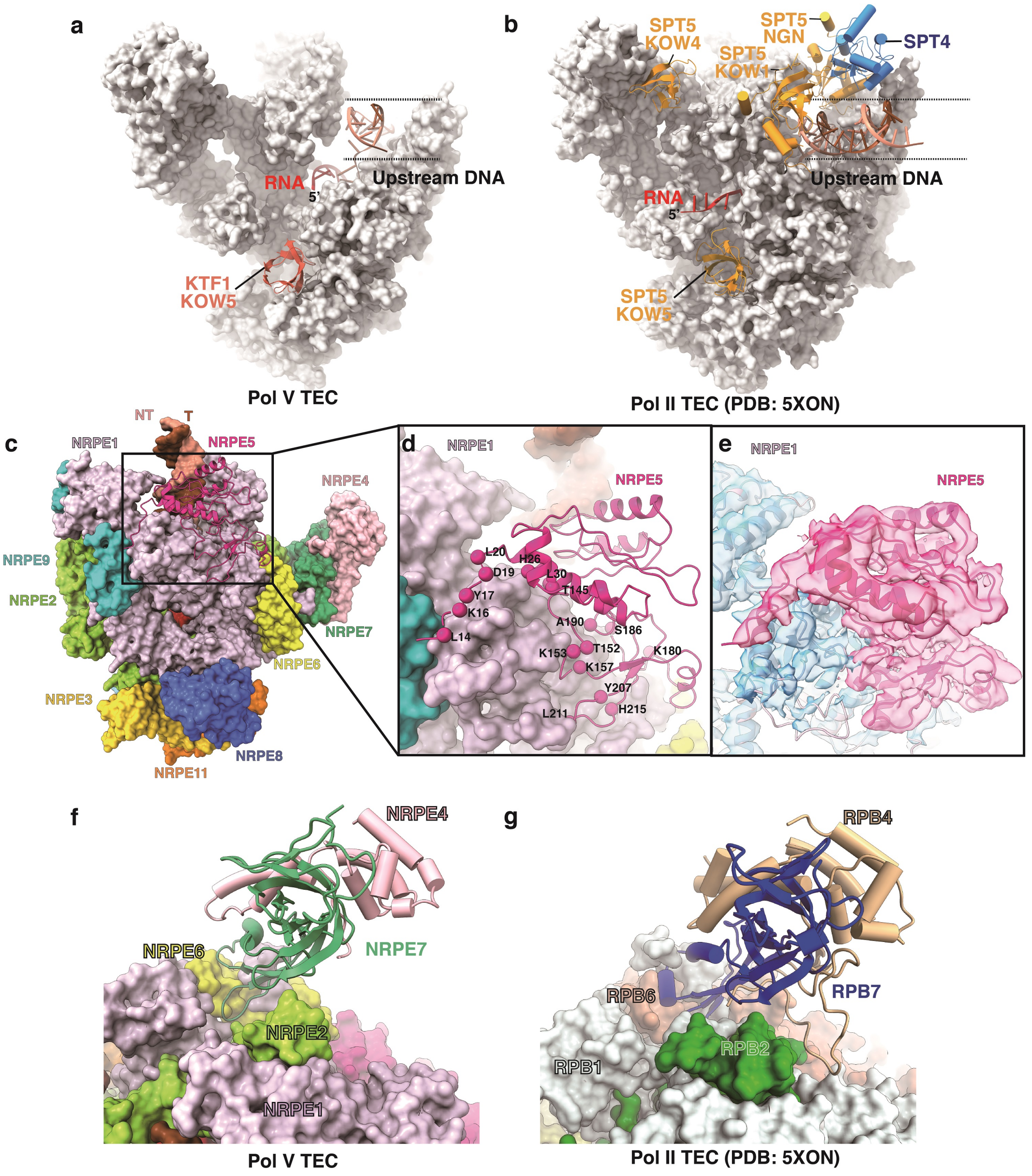


**Supplementary Figure 5. The interactions of Pol V-specific subunits and recruting factror KTF1. (a)** The interaction between KTF1 KOW5 (pink) and Pol V. **(b)** The interaction between SPT5-SPT4 heterodimer (yellow and blue) and Pol II (PDB: 5XON). **(c)** The overall structure of Pol V TEC shows the interaction between NRPE1 (surface) and NRPE5 (cartoon). **(d)** Close-up view of the NRPE1-NRPE5 interface. Red spheres indicate the NRPE5 residues that are invovled in the interaction with NRPE1 and not conserved in NRP(A/B/C/D)5. **(e)** The local cryo-EM map (map 4) for the NRPE5 subunit and the cryo-EM map (map 2) for the NRPE1 subunit. **(f)** The interface between NRPE1-NRPE7 in the Pol V TEC. **(g)** The interface between RPB1-RPB7 in the Pol II TEC (PDB: 5XON).

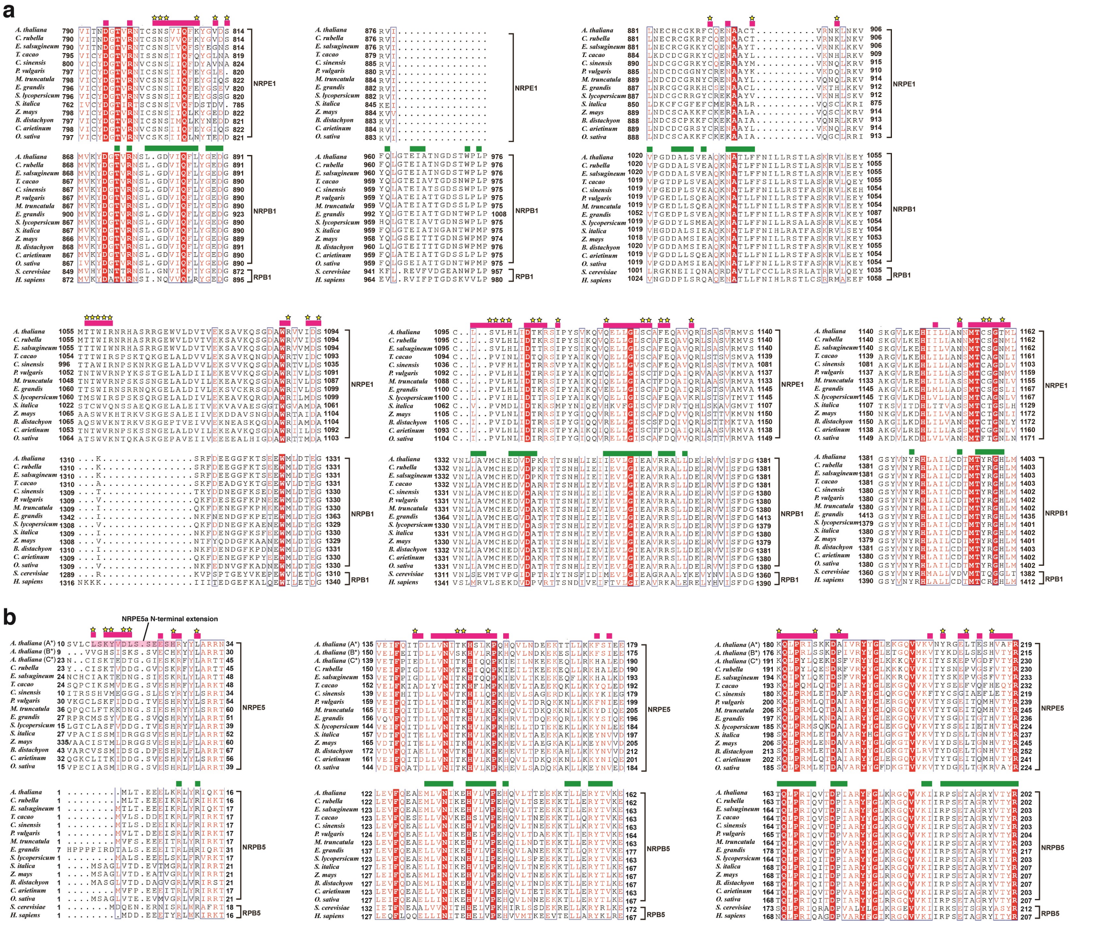


**Supplementary Figure 6. The sequence alignment of Pol II, Pol IV, and Pol IV showing interface residues on (a) NRPE1/RPB1 and (b) NRPE5/RPB5.** The pink bars label Pol V residues that participate into the interactions between NRPE1 and NRPE5. The green bars label Pol II residues that participate into the interactions between RPB1 and RPB5. Stars represent interface residues nonconserved between Pol II and Pol V. The NRPE5-specific N-terminal extension is highlighted by a transparent pink box in the NRPE5a sequence. Our LC-MS/MS result shows the presence of both NRPE5a (At3g57080) and NRPE5c (At3g54490) in the epitope-tagged Pol V, in contrast to the presence of only NRPE5a in the previous study[2]. The discrepancy likely results from the experimental materials used for Pol V immunoprecipitation.

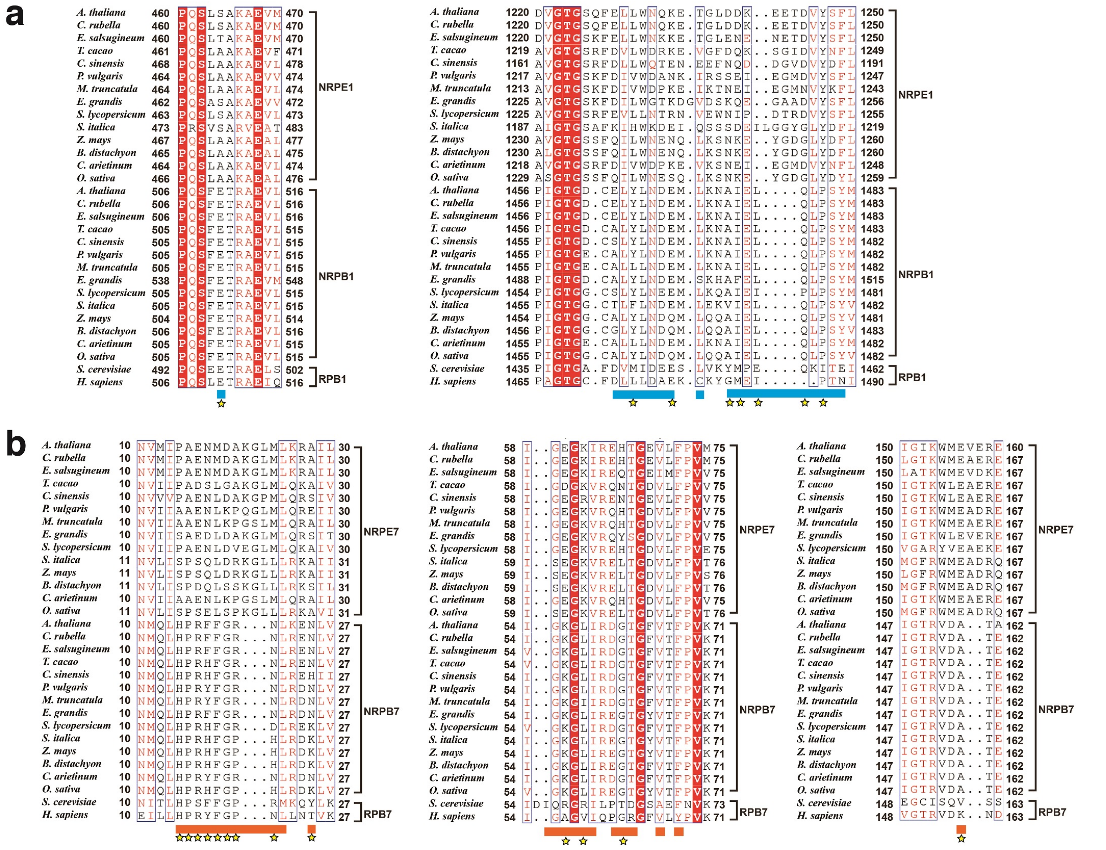

**Supplementary Figure 7. The sequence alignment of Pol II, Pol IV, and Pol IV showing interface residues on NRPE1/RPB1 (a) and NRPE7/RPB7 (b).** The yellow bars label Pol II residues that participate into the interactions between RPB1 and RPB7. Stars represent interface residues nonconserved between Pol II and Pol V.

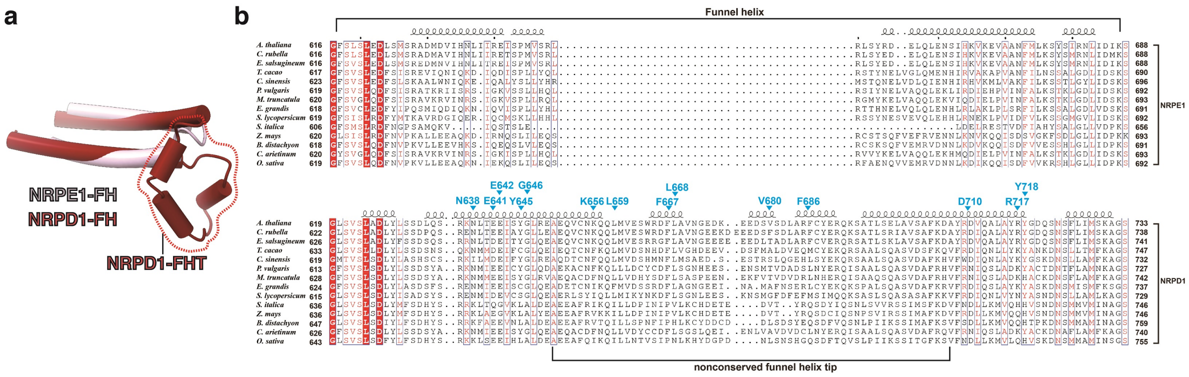


**Supplementary Figure 8. The sequence alignment of the Pol IV-NRPD1 subunit and the Pol V-NRPE1 subunit showing the differences of the funnel helices. (a)** The structural superimpostion of NRPE1-FH and NRPD1-FH (PDB: 7EU0)[3].The Pol IV-specific insertion of at the funnel helix tip (NRPD1-FHT) is highlighted by dashes. **(b)** Pol IV-NRPD1 residues making interactions with RDR2 are labeled.

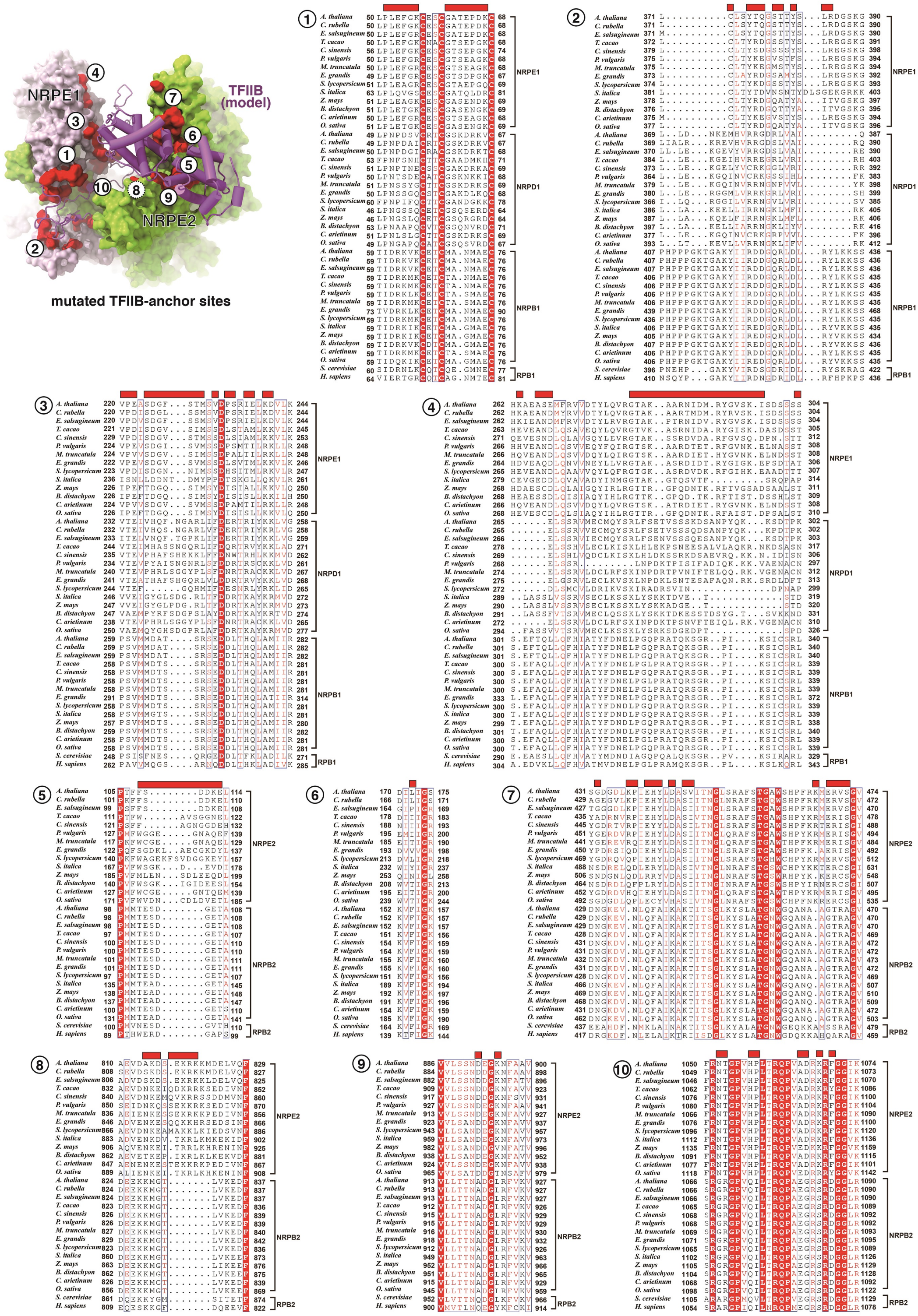


**Supplementary Figure 9. Pol V is incompatible with TFIIB interaction.** The red surface patches highlight non-conserved residues of Pol V on the corresponding TFIIB-contact surface of Pol II. The red bars highlight non-conserved residues of Pol V on the corresponding TFIIB-contact surface of Pol II.

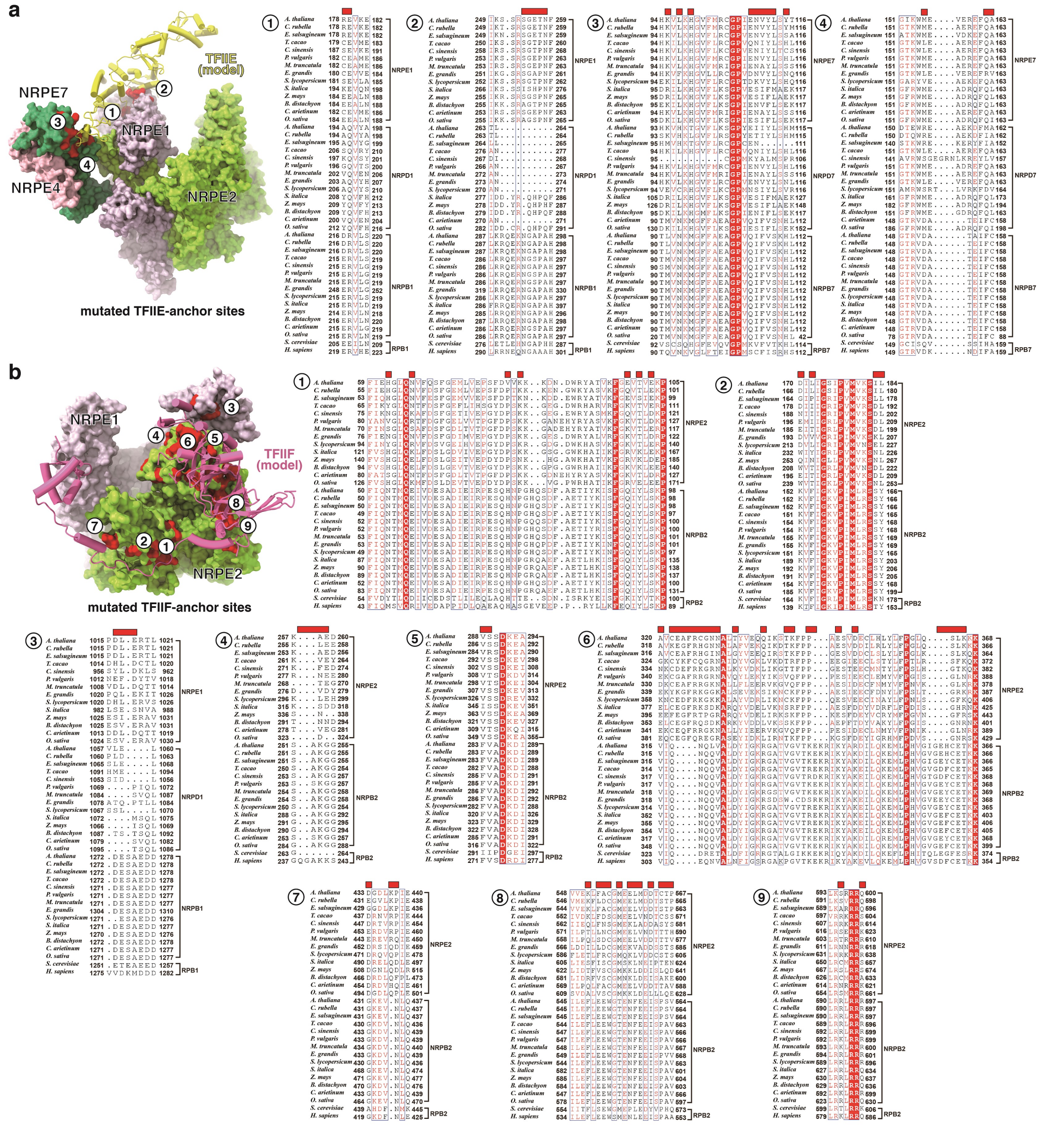


**Supplementary Figure 10. Pol V is incompatible with the interactions with (a) TFIIE and (b) TFIIF.** The red surface patches highlight non-conserved residues of Pol V on the corresponding TFIIE/F-contact surface of Pol II. The red bars highlight non-conserved residues of Pol V on the corresponding TFIIE/F-contact surface of Pol II.

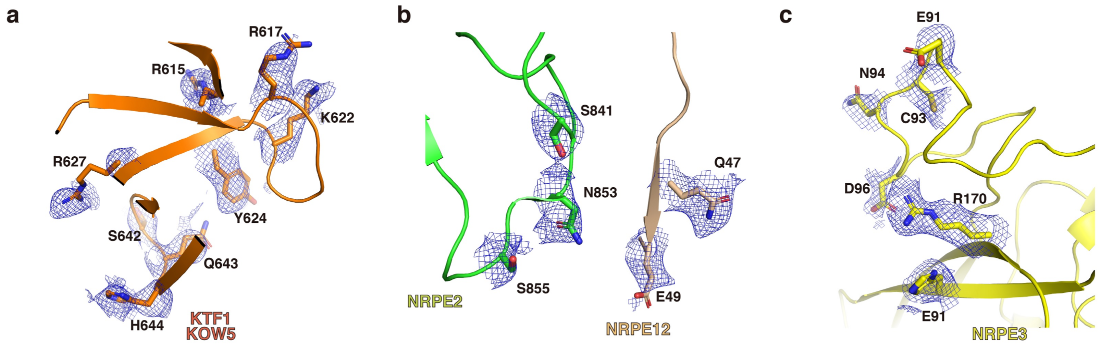


**Supplementary Figure 11. The cryo-EM map (blue dash; map 3) for the interface residues of Pol V and KTF1.**

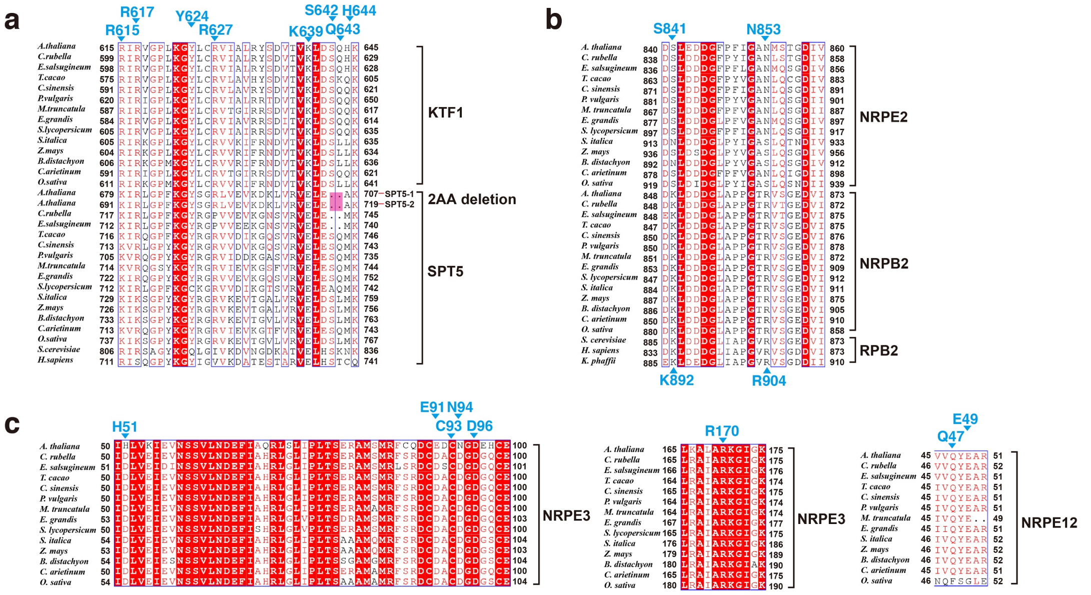


**Supplementary Figure 12. Sequence alignments showing shared or non-conserved interfaces of the KTF1-Pol V interaction and SPT5-Pol II interaction. (a)** The sequence alignment of KTF1 and SPT5. **(b)** The sequence alignment of the NRPE2 subunit of Pol V and NRPB2 subunit of Pol II. **(c)** The sequence alignment of NRP (B/D/E)3 and NRP (A/B/C/D/E)12 subunits. The interface residues participating in interactions between the KTF1 KOW5 domain and Pol V are labeled.

**Supplementary Table 1. The subunit composition of Arabidopsis Pol II, IV and V[2].**

| Subunit | Pol II | Pol IV | Pol V |
|---|---|---|---|
| 1 | NRPB1 | NRPD1 | NRPE1 |
| 2 | NRPB2 | NRP(D/E)2 | |
| 3 | NRP(B/D/E)3 | | |
| 4 | NRPB4 | NRP(D/E)4 | |
| 5 | NRP(A/B/C/D)5 | | NRPE5 |
| 6 | NRP(A/B/C/D/E)6 | | |
| 7 | NRPB7 | NRPD7 | NRPE7 |
| 8 | NRP(A/B/C/D/E)8 | | |
| 9 | NRP(B/D/E)9 | | |
| 10 | NRP(A/B/C/D/E)10 | | |
| 11 | NRP(B/D/E)11 | | |
| 12 | NRP(A/B/C/D/E)12 | | |

**Supplementary Table 2. Pol V subunits detected by LC-MS/MS of affinity purified epitope-tagged Pol V from *At* T87 cells.**

| AGI code | Pol V subunit | MW (kDa) | Peptides | Unique peptides* | Coverage (%) |
|---|---|---|---|---|---|
| AT2G40030 | NRPE1 | 218.2 | 75 | 65 | 38% |
| AT3G23780 | NRP(D/E)2 | 132.7 | 53 | 43 | 38% |
| AT2G15400 | NRP(B/D/E)3b | 35.5 | 7 | 7 | 28% |
| AT4G15950 | NRP(D/E)4 | 22.4 | - | - | - |
| AT3G57080 | NRPE5a | 25.6 | 11 | 9 | 32% |
| AT3G54490 | NRPE5c | 26.9 | 6 | 5 | 19% |
| AT5G51940 | NRP(A/B/C/D/E)6a | 16.7 | - | - | - |
| AT4G14660 | NRPE7 | 20.3 | 6 | 6 | 37% |
| AT1G54250 | NRP(A/B/C/D/E)8a | 16.5 | 2 | 1 | 9% |
| AT3G16980 | NRP(B/D/E)9a | 13.3 | 7 | 4 | 44% |
| AT4G16265 | NRP(B/D/E)9b | 12.3 | 2 | 2 | 37% |
| AT1G11475 | NRP(A/B/C/D/E)10 | 8.3 | 3 | 2 | 28% |
| AT3G52090 | NRP(B/D/E)11 | 13.6 | 6 | 5 | 52% |
| AT5G41010 | NRP(A/B/C/D/E)12 | 5.9 | - | - | - |

* unique peptide: the peptide sequence that is unique to a certain proteome and can specifically distinguish it from other proteomes.
- no peptides were identifed.

**Supplementary Table 3: The statistics of cryo-EM structure determination.**

| | Pol V TEC Map 1 | Pol V TEC Map 2 | Pol V TEC Map 3 | Pol V TEC Map 4 |
|---|---|---|---|---|
| **Data collection and Processing** | | | | |
| Microscope | Titan Krios | | | |
| Voltage (kV) | 300 | | | |
| Camera | Gatan K3 | | | |
| Magnification | 64000 | | | |
| Pixel size at detector (Å/pixel) | 1.1 | | | |
| Total electron exposure ($e^{-}$/Å$^2$) | 50 | | | |
| Exposure rate (e-/pixel/sec) | 22.5 | | | |
| Number of frames | 40 | | | |
| Defocus range (μm) | -1.2 to -2.2 | | | |
| Micrographs collected (no.) | 6811 | | | |
| Micrographs used (no.) | 6811 | | | |
| Total extracted particles (no.) | 3436012 | | | |
| **<u>For each reconstruction:</u>** | | | | |
| Refined particles (no.) | 30359 | 273007 | 273007 | 273007 |
| Final particles (no.) | 30359 | 273007 | 273007 | 273007 |
| Point-group or helical symmetry parameters | C1 | C1 | C1 | C1 |
| Resolution (global, Å) | | | | |
| FSC 0.143 (unmasked/masked) | 4.2/4.7 | 3.2/3.5 | 3.1/3.3 | 3.9/4.0 |
| Resolution range (local, Å) | 3.7-10.0 | 3.0-10.0 | | |
| Map sharpening *B* factor (Å$^2$) | -120.1 | -90.3 | -78.1 | -80.6 |
| **Model composition** | | | | |
| Protein | 3529 | | | |
| Ligands | 4/1 | | | |
| RNA/DNA | 83 | | | |
| **Model Refinement** | | | | |
| resolution cutoff | 4.26 | | | |
| Model-Map scores | | | | |
| CC | 0.52 | | | |
| *B* factors (Å$^2$) | | | | |
| Protein residues | 222.9 | | | |
| Ligands | 195.8 | | | |
| RNA/DNA | 477.9 | | | |
| R.m.s. deviations from ideal values | | | | |
| Bond lengths (Å) | 0.027 | | | |
| Bond angles (°) | 2.35 | | | |
| **Validation** | | | | |
| MolProbity score | 2.36 | | | |
| CaBLAM outliers | 3.64 | | | |
| Clashscore | 11.5 | | | |
| Poor rotamers (%) | 3.24 | | | |
| C-beta deviations | 8.12 | | | |
| Ramachandran plot | | | | |
| Favored/Allowed/Outliers (%) | 94.1/5.9/0 | | | |

**Supplementary references**


1 Ehara, H. *et al.* Structure of the complete elongation complex of RNA polymerase II with basal factors. *Science* **357**, 921-924, doi:10.1126/science.aan8552 (2017).

2 Ream, T. S. *et al.* Subunit compositions of the RNA-silencing enzymes Pol IV and Pol V reveal their origins as specialized forms of RNA polymerase II. *Mol Cell* **33**, 192-203, doi:10.1016/j.molcel.2008.12.015 (2009).

3 Huang, K. *et al.* Pol IV and RDR2: A two-RNA-polymerase machine that produces double-stranded RNA. *Science* **374**, 1579-1586, doi:10.1126/science.abj9184 (2021).
